# Species level enrichment of AnAOB and associated growth morphology under the effect of key metabolites

**DOI:** 10.1101/2020.02.04.934877

**Authors:** Yang Lu, Gayathri Natarajan, Thi Quynh Ngoc Nguyen, Sara Swa Thi, Krithika Arumugam, Thomas William Seviour, Rohan B.H. Williams, Stefan Wuertz, Yingyu Law

**Affiliations:** Singapore Centre for Environmental Life Sciences Engineering, Nanyang Technological University, Singapore 637551, Singapore; Singapore Centre for Environmental Life Sciences Engineering, National University of Singapore, Singapore 119077, Singapore; School of Civil and Environmental Engineering, Nanyang Technological University, Singapore. 639798, Singapore

**Keywords:** *Candidatus* Brocadia, Granule, Nitrogen load, *Candidatus* Jettenia, Nitric oxide, Organic carbon

## Abstract

Nineteen anaerobic ammonium oxidizing bacteria (AnAOB) species have been identified, yet the environmental factors that select for each species within a specialized ecological niche have not been described. We enriched AnAOB from a single inoculum under standard enrichment conditions (reactor R1) with stepwise increase in nitrite and ammonia concentration, nitric oxide (NO) supplementation (reactor R2), or with complex organic carbon using wastewater collected from mainstream wastewater treatment facility (reactor R3). AnAOB were enriched up to 80%, 90% and 50% relative abundance in R1, R2 and R3 respectively. *Candidatus* Brocadia caroliniensis predominated in all reactors, but a shift towards *Ca*. Brocadia sinica was consistently observed with increasing ammonium and nitrite concentrations beyond 270 mg NH_4_-N L^−1^ and 340 mg NO_2_-N L^−1^, respectively. In the presence of NO, growth of heterotrophs were inhibited, and *Ca*. Jettenia could coexist with *Ca*. B. caroliniensis before diminishing when nitrite increased to 160 mg NO_2_-N L^−1^. In contrast, supplementation of organic carbon led to the emergence of heterotrophic communities that coevolved with *Ca*. B. caroliniensis. *Ca*. B. caroliniensis and *Ca*. Jettenia preferentially form biofilms on reactor surfaces, whereas *Ca*. Brocadia sinica forms granules in suspension. Our results thus indicate that multiple AnAOB species co-exist and occupy sub-niches in anaerobic ammonium oxidation reactors, that the dominant population can be reversibly shifted by, for example, changing the nitrogen load (i.e. high nitrite concentration favors *Ca*. Brocadia caroliniensis), and that speciation has implications for wastewater process design, with the optimum cell immobilization strategy (i.e. carriers vs granules) dependent on which species dominates.

**IMPORTANCE:** This study demonstrates how to reversibly and predictably shift dominant anammox population using operating parameters (e.g. high nitrite concentration favours *Ca*. Brocadia sinica), and that species selection has implications for wastewater process design, illustrated here in terms of dependence of optimum cell immobilization strategy (i.e. carriers vs granules) on which species dominates. The research informs the characterization of AnAOBs at species level as well process design and control strategies targeting Anammox species population dynamics in full scale waste water treatment systems.

## INTRODUCTION

Anaerobic ammonium oxidation (Anammox), combined with partial nitritation is applied widely to treat nitrogen-rich and carbon-deficient wastewaters (e.g., sidestream treatment) due to significant energy savings relative to conventional processes. It has also been proposed as a sustainable treatment option for treating municipal wastewaters (i.e. mainstream treatment). Nineteen anaerobic ammonium oxidizing bacteria (AnAOB) species have been identified in various environments, including suboxic marine zones, coastal sediments, lakes, and wastewater treatment plants. These have been classified into five candidatus genera **(1-3)**, and while AnAOB can colonize diverse natural and engineered systems, different genera rarely coexist in the same habitat **(4)**. Differences in growth rates, substrate affinities, sensitivities to inhibitory compounds, preferred growth substrates and differential metabolic pathways, are all thought to contribute to niche specialization **(1, 5-10)**.

However, population shifts at species and genus level have been reported in Anammox lab-scale reactors under various conditions **(9-11)**. In fact, during scale-up of the first full-scale commercial Anammox reactor, the dominant population shifted from *Ca*. Kunenia stuttgartiensis to *Ca*. Brocadia anammoxidans, although reasons for this were not provided **(12)**. Studies have reported that specific environmental conditions in partial nitritation/Anammox (PN/A) reactors can select for single AnAOB species only **(11, 13)**. For instance, *Ca*. Jettenia moscovienalis **(2)**, *Ca*. B. caroliniensis **(14)**, and *Ca*. B. sinica **(13)** were detected in distinct sidestream reactors treating anaerobic digester liquor, whereas *Ca*. B. sp. 40 was identified as the dominant AnAOB under mainstream conditions **(15)**. Park et al. **(11)** showed that feed composition is more important in AnAOB selection than inoculum and reactor configuration. Nonetheless, there is no apparent consensus on which factors select for one AnAOB species over another.

Elucidating factors that enrich for specific AnAOB species with specific kinetic and physiological properties would potentially enhance process design and performance. AnAOB exist in a range of conditions, such as sidestream and mainstream PN/A systems with high and low ammonium/nitrite concentrations respectively, and many factors are likely involved in species selection. Nitrite (NO_2_^−^) acts as an electron acceptor for ammonium oxidation and electron donor for bicarbonate reduction to biomass. It applies an AnAOB-species selection pressure on the basis of their abilities to utilize it, but also tolerate it as NO_2_^−^ can be inhibitory to their growth. A 50 % reduction in Anammox activity has been reported in AnAOB exposed to concentrations between 100 and 400 mg N L^−1^ **(16-18)**. Nitric oxide (NO), a potent oxidant produced from nitrite as an important intermediate in the Anammox biochemical pathway **(19, 20)**, is toxic to many bacteria; yet AnAOB can tolerate it at higher concentrations **(21, 22)**. In addition to potential selection pressures from nitrite and NO, the ability to also consume organic substrates (i.e. acetate and propionate) conferred competitive advantages to *Ca*. B. fulgida and *Ca*. Anammoxoglobus propionicus respectively, over other species including denitrifers **(6, 7)**. It remains to be determined whether selection of such ‘facultative chemoorganotrophs’ would be favored in the complex organic carbon milieu present under mainstream conditions. Nonetheless, the AnAOB occupy highly specialized niches that are defined by more than just the concentrations of ammonium and nitrite.

The ability to enhance the activity of specific AnAOB species with favorable physiological and growth properties is particularly advantageous when starting up and optimizing industrial anammox processes. Inoculating from existing full-scale installations to start up Anammox reactors is logistically challenging **(11)** or simply unfeasible owing to the sensitivity of the process to feed composition, oxygen **(12)** and competing microbial species **(23)**. High biomass retention is required in Anammox reactors due to the slow growth rates of AnAOB. This can be achieved by promoting biomass aggregation in biofilm-based Anammox reactors **(24)**.

Biomass in an Anammox reactor can self-assemble into flocs in suspension, fixed films on surfaces or carriers, small granules, big granules or some combination of all of these morphologies **(25)**. Such aggregates can play functionally different roles within the reactor and even affect nitrogen removal efficiencies **(26, 27)**. Understanding which factors drive species selection and whether different species assume particular biofilm morphologies could inform process design and control strategies for achieving more stable nitrogen removal by Anammox.

This work aims to describe how Anammox community composition, process performance and biofilm morphology are shifted by factors typically encountered in industrial Anammox systems (i.e. mainstream compared to sidestream). We hypothesized that different substrate compositions, simulating a) organic carbon-deficient wastewaters with N-loads spanning domestic mainstream and sidestream wastewaters (reactors R1) and with b) an oxidative stress typically encountered in PN/A systems through the exposure to NO (reactor R2), or c) domestic strength mainstream wastewater with high COD:N (reactor R3), could select for distinct community, specifically AnAOB species. While some of these factors have been previously investigated independently **(17, 28)**, this study systematically investigates these factorson AnAOB species selection from the same inoculum. Resolving the factors selecting Anammox species is important for N-removal process design, and may contribute to the understanding of niche partitioning in complex microbial/environmental habitats.

## RESUTS

### Start-up period reduced and N removal activity enhanced with NO supplementation

AnAOB were successfully enriched under all tested enrichment conditions albeit with varying start-up times. Start-up period was the shortest in R2 supplemented with NO and Anammox activity was observed within 20 days of inoculation compared to 39 days for R1, operated under standard enrichment conditions (Figs 1A & B). In the presence of complex organic carbon in R3, Anammox activity was only detected after 50 days of operation (Fig 1C). In both R1 and R2, ammonium and nitrite concentrations were increased to 280 mg N L^−1^ and 350 mg N L^−1^, respectively (Fig 1A and B), above which anammox activity was inhibited. In addition to the shorter start-up period, a shorter HRT was applied to R2 than R1 due to higher N removal rates of 1200 mg N L^−1^ day^−1^ compared to 800 mg N L^−1^ day^−1^ under stable operation (Fig 1A and B). Despite the higher loading rate, the suspended solid concentrations were comparable in both reactors indicating a higher specific N removal activity for R2 than R1. A significantly lower N loading rate of 121 ± 6 mg N L^−1^ day^−1^ was achieved in R3. Final effluent ammonium concentration dropped steadily from day 58 to 80, with the reactor displaying stable ammonium removal activity by AnAOB henceforth. Residual nitrite in the effluent also decreased gradually from day 60 to 100 along with a decrease in ammonium concentration (Fig 1C). The average TCOD and sCOD in the effluent were 87±9 mg L^−1^ and 51±8 mg L^−1^, respectively with an average removal rate of 520 mg L^−1^ day^−1^ from day 300 onwards.

**Figure 1.**
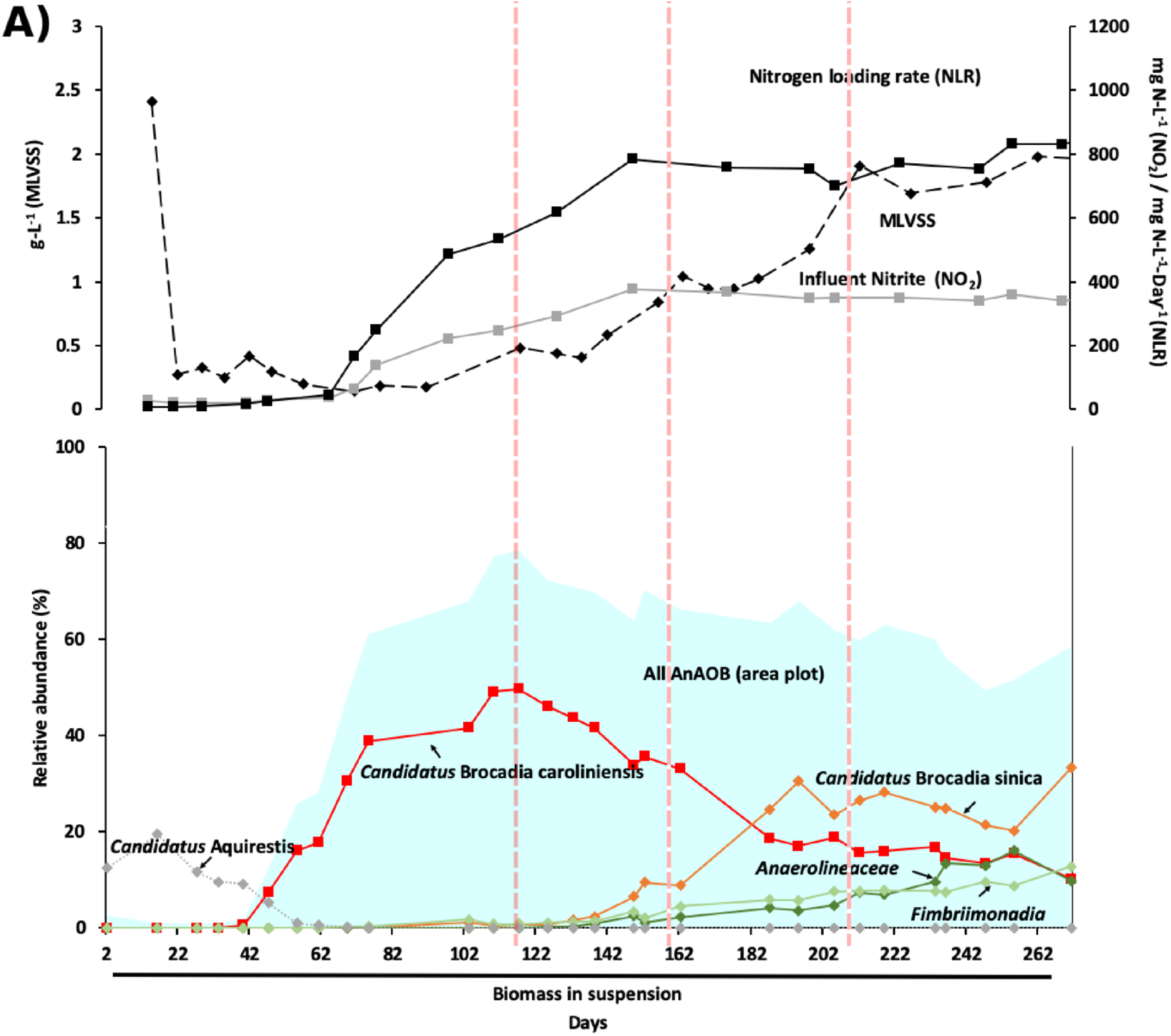
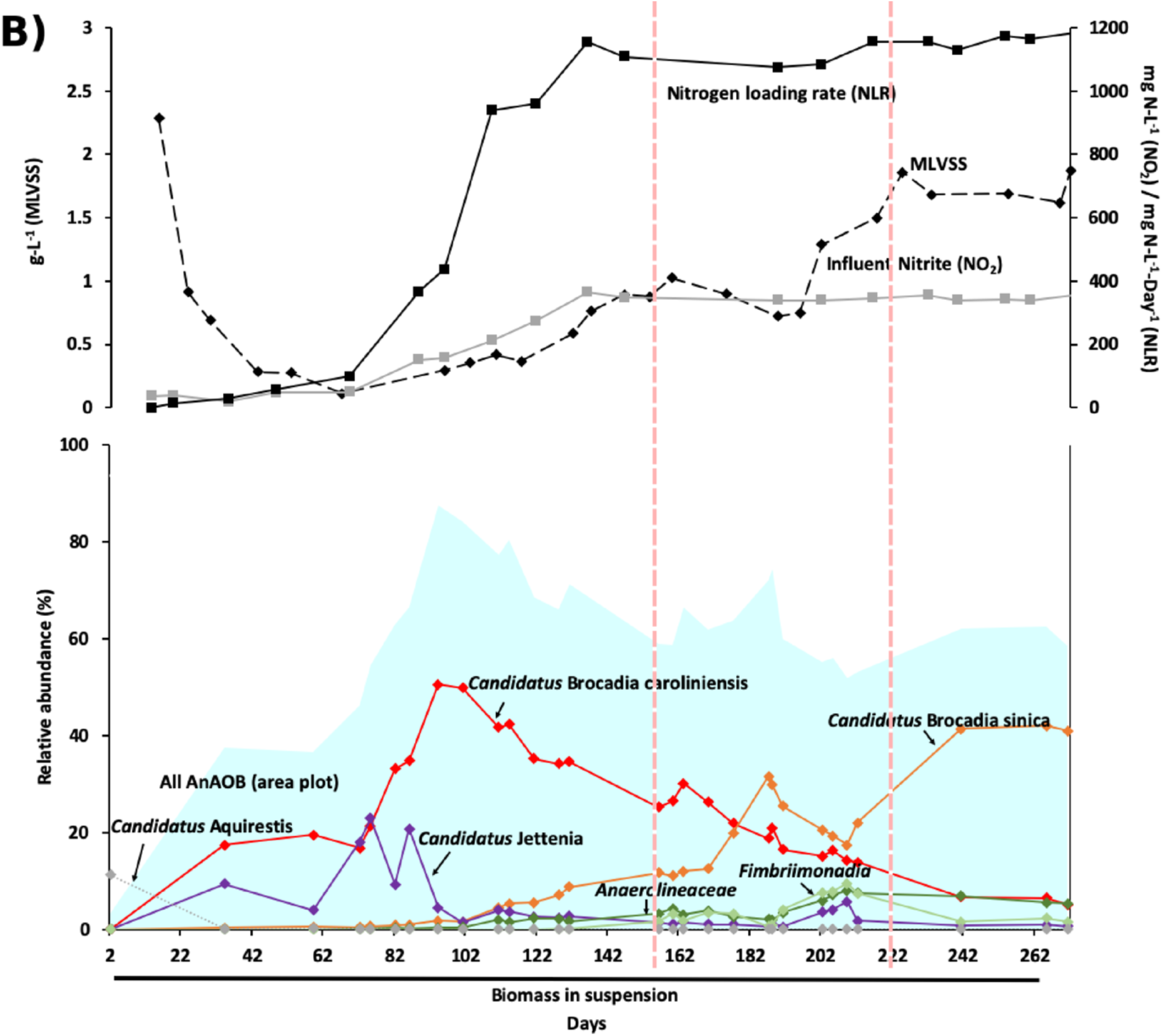
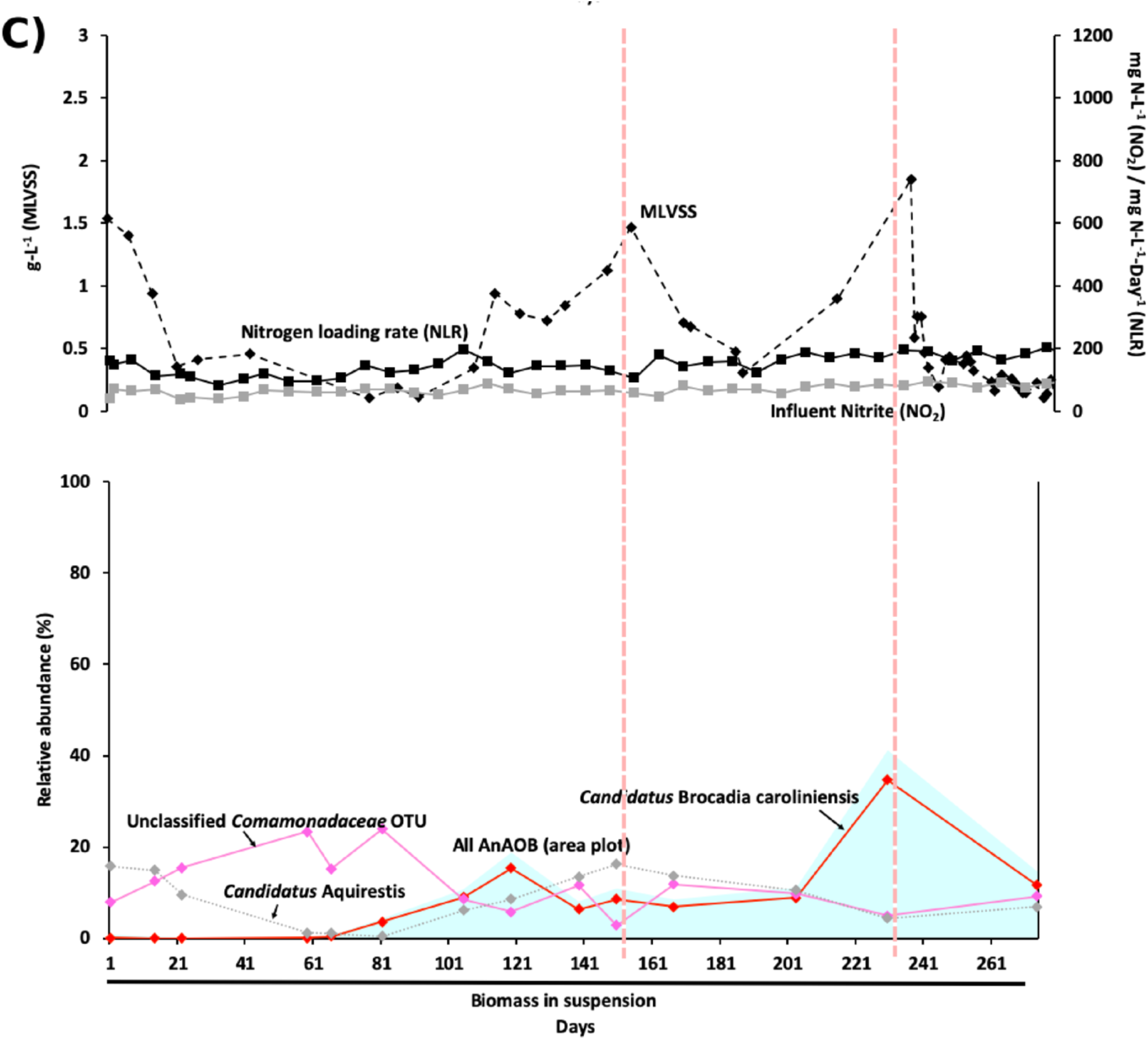
Start-up and enrichment of AnAOB from activated sludge fed with (A) synthetic waste water with ammonium and nitrite in <u>R1</u>, (B) synthetic waste water with ammonium, nitrite and continuous supply of nitric oxide in <u>R2</u>, and (C) primary effluent supplemented with nitrite in <u>R3</u>. Mixed liquid volatile suspended solids (MLVSS, 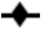), Influent nitrite (NO_2_, 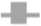) and nitrogen loading rates (NLR, 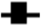) of each enrichment condition are shown in the upper panel of each graph; with relative abundance of dominant AnAOB OTUs affiliated to *Ca*. B. caroliniensis 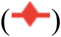 and *Ca*. B. sinica 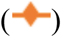 and correlated non-AnAOB OTUs affiliated to *Anaerolineaceae* 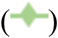 and *Fimbriimonadia* 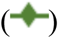 are shown in the lower panels of A and B; *Ca*. Jettenia 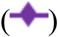 was also detected in <u>R2</u> (B). *Ca*. B. caroliniensis, the only dominant AnAOB in <u>R3</u>, is shown in C along with correlated non-AnAOB OTUs affiliated to *Comamonadaceae* 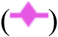 and *Ca*. Aquirestis 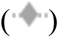 dominating at a different stage. The relative abundance of total AnAOB is highlighted as area plot 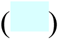 in each graph. Red dotted lines denote the time points at which Anammox biofilm was scraped from the wall of the reactor into suspension. The detailed chemical and microbial community (with OTUs >5% at any analysed time point) can be found in Figure S1, Supporting Information.

### High N load shifts the dominant AnAOB from *Ca*. B. caroliniensis to *Ca*. B. sinica

Along with the increase in Anammox activity, a shift in the functional AnAOB was observed in R1 and R2 along with increasing N load, but not in R3 where low N load was maintained. 16S rRNA gene amplicon sequencing showed that AnAOB were below the detection limit at the start of reactor operation for all three reactors. In R1 and R2, OTUs annotated to AnAOB increased progressively to 80% (day 110) and 90% (day 95) relative abundances, respectively, at influent nitrite concentration > 200 mg N L^−1^ (Fig 1A and B). Despite the proliferation of multiple OTUs affiliated to AnAOB in both R1 and R2, a single OTU annotated to *Ca*. Brocadia, identified as *Ca*. B. caroliniensis by clone library analysis (Fig 2), dominated throughout the first 120 days of reactor operation. *Ca*. B. caroliniensis increased during enrichment to 50% relative abundance in R1 and R2. However, a further increase in influent ammonium and nitrite concentrations beyond 220 mg N L^−1^ from day 100 (N loading rate of 500 mg N L^−1^-day^−1^ for R1 and 750 mg N L^−1^-day^−1^ for R2) resulted in the gradual increase of *Ca*. Brocadia_2, identified as *Ca*. B. sinica by clone library analysis (Fig 2).

**Figure 2.**
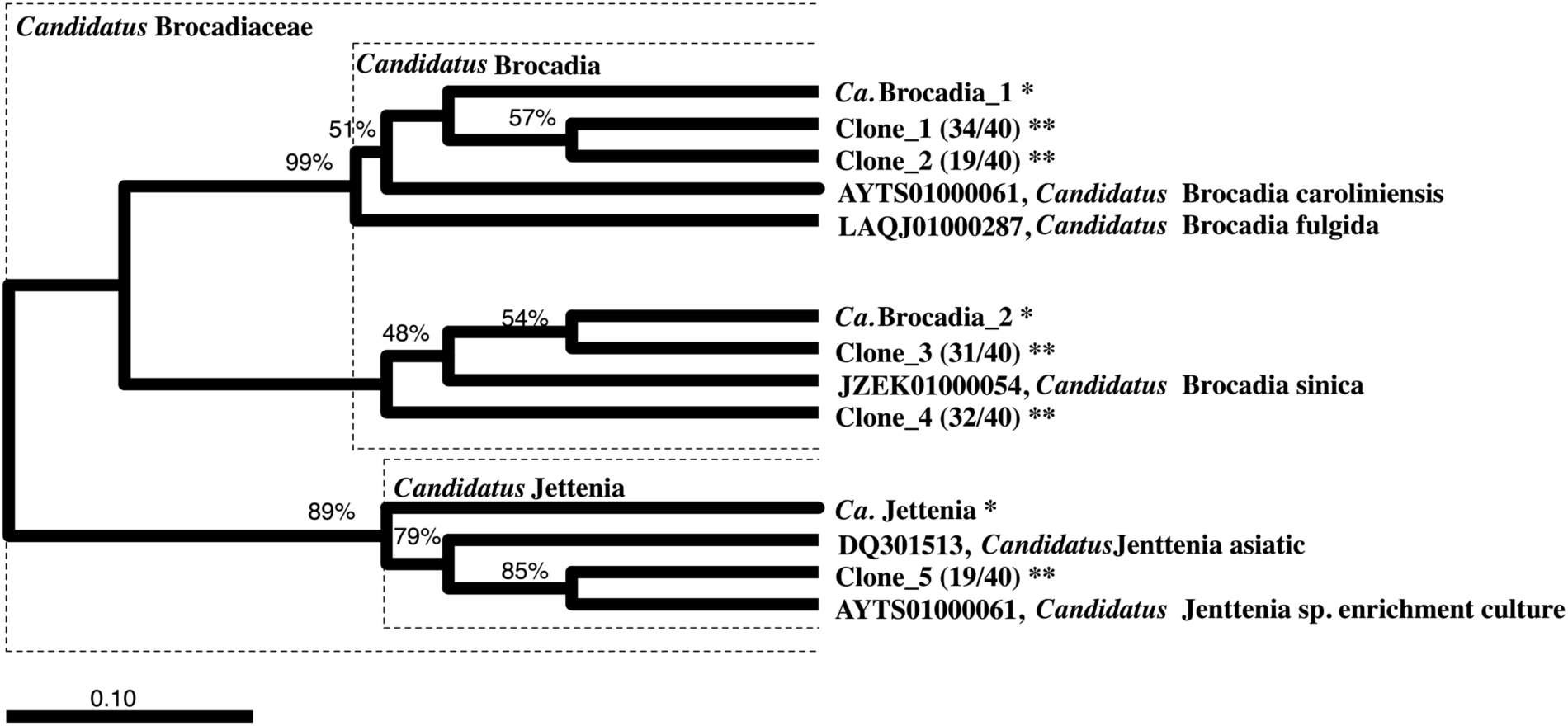
Phylogenetic tree based on 16S rRNA sequences of major OTUs (with postfix “*”) and clones (with postfix “**”) from amplicon sequencing and clone library analysis, respectively. Number of identical colonies per total colonies picked is indicated in parentheses followed by reactor and day for sample collection. The phylogenetic tree was generated by ARB with SILVA database. Sequences obtained from clone library and amplicon sequencing were inserted into the tree using parsimony insertion tool of ARB. The closest neighbor sequences were selected to generate the final tree with neighbor-joining method with bootstrap of 1000 replications. Dashed line indicates the division of family *Ca*. Brocadiaceae and genus *Ca*. Brocadia. *Ca*. Kuenenia stuttgartiensis was selected as the root (not shown). Only the closest identified sequences were selected to be shown. The scale indicates 0.1 nucleotide change per nucleotide position. Sequence affiliations and relative abundances of major AnAOB were consistent between amplicon sequencing and clone library analysis.

A decrease in *Ca*. B. caroliniensis was also observed (Fig 1A & B). Beyond 180 days of reactor operation, *Ca*. B. sinica increased to 33% and 42% in relative abundance in R1 and R2, respectively, while *Ca*. B. caroliniensis decreased to less than 10% in relative abundance in both reactors. In the presence of COD, *Ca*. B. caroliniensis was the most dominant Anammox taxon throughout the enrichment process in R3 when operated at a low N loading rate of 120 mg N L^−1^ day^−1^ (Figs 1C). However, the relative abundance of total AnAOB was significantly lower in R3 (∼50%) than in R1 and R2 (∼80%), suggesting a more competitive environment for AnAOB in the presence of organic carbon. FISH analysis (Fig 3D, E and F) on R1 (day 80) and R2 (day 675) further indicated that *Ca*. B. sinica dominated in those reactors while *Ca*. B. caroliniensis remained as the only AnAOB detected in R3 (day 683). The designed species-specific FISH probes served to observe gradual population shifts during reactor operation in response to changes in the controlling factors.

**Figure 3.**
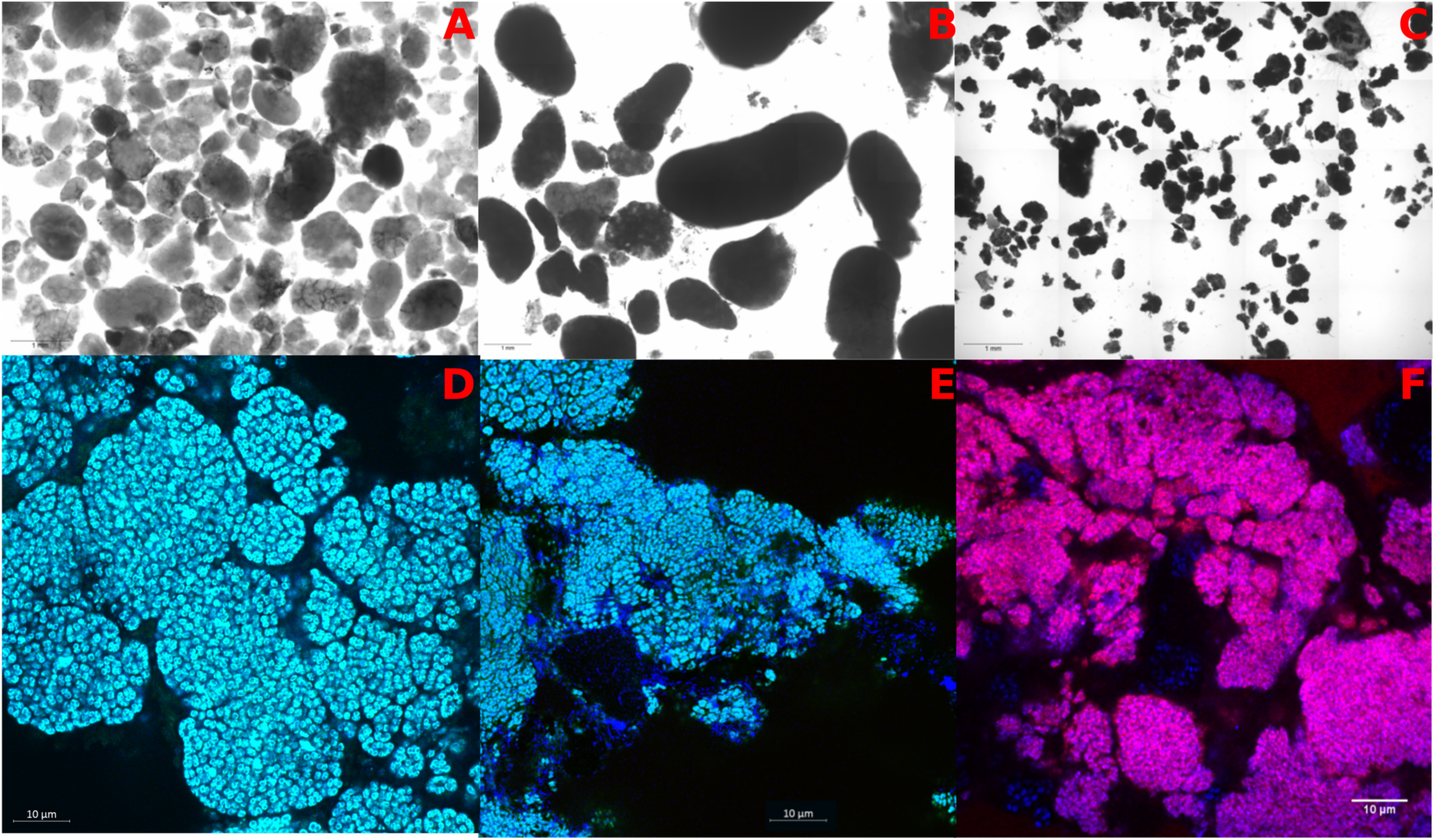
Light microscopy images of suspended biomass samples showing granular structure in R1 (A) and R2 (B) and floccular structure in R3 (C). FISH images of crushed granules were taken with *Ca*. B. sinica (cyan), *Ca*. B. caroliniensis (magenta) and other AnAOB (blue) for R1 (D) R2 (E) and R3 (F) at the end of enrichment.

While other OTUs affiliated to genus *Ca*. Brocadia were also detected, their relative abundance was less than 10% (Fig S1). The presence of these *Ca*. Brocadia OTUs is likely due to different strains of *Ca*. Brocadia or sequencing errors as only low relative abundances were detected. Aside from *Ca*. Brocadia, OTUs affiliated to *Ca*. Jettenia emerged to coexist with *Ca*. B. caroliniensis only in R2 (with continuous supply of NO), suggesting that the presence of NO may provide a competitive advantage for *Ca*. Jettenia. However, *Ca*. Jettenia diminished with increasing N loading rate (i.e. after day 102).

### Excess NO availability over nitrite selects for *Ca*. Jettenia

*Ca*. Jettenia was only observed in R2 but not in R1 and R3. We posit that this was due to the presence of NO as it was the only condition that differed between R1 and R2. However, during the enrichment process it was unclear whether the NO effect was due to imposition of an oxidative stress or because it was used as a substrate for ammonium oxidation. This could not be assessed because nitrite was in excess during the enrichment. Hence, to investigate whether NO is consumed, nitrite was systematically depleted in R2 (Phase II) while dosing the same amount of NO (Phase I). After 76 days of nitrite depletion, nitrite was gradually reintroduced in Phase III (Fig 4).

**Figure 4.**
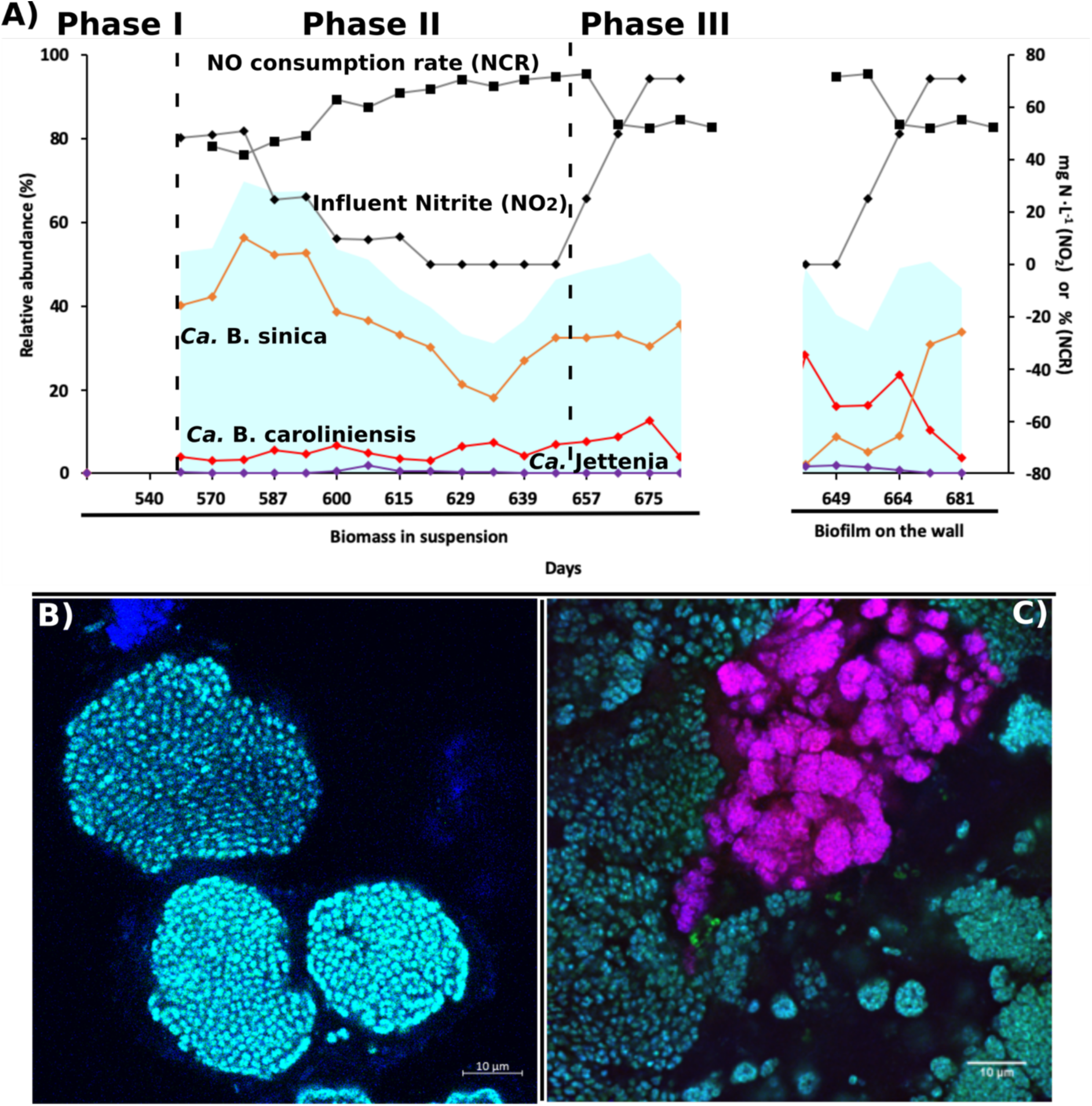
The effect of nitrite depletion (Phase II) and repletion (Phase III) on the changes in (A) AnAOB community of OTUs affiliated to *Ca*. B. caroliniensis 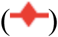 and *Ca*. B. sinica 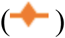 and *Ca*. Jettenia 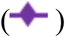 in suspended (highlighted as “Granules in suspension” on the x-axis) and attached growth biomass (highlighted as “biofilm on the wall” on the x-axis), and the NO consumption rate (NCR, 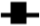) of R2, fed with synthetic waste water with ammonium, nitrite and continuous supply of nitric oxide. Influent nitrite (NO_2,_ 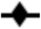) was adjusted from normal (Phase I), to depletion (Phase II) and repletion (Phase III). The relative abundance of total AnAOB is highlighted as area plot 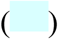. FISH images were taken with *Ca*. B. sinica in cyan and *Ca*. B. caroliniensis in magenta during Phase I (B) and Phase III (C) from crushed granules.

Reducing the nitrite concentration resulted in a decrease in the relative abundance of *Ca*. B. sinica in suspension while a slight increase was observed for both *Ca*. B. caroliniensis and *Ca*. Jettenia in Phase II (Fig 4). The presence of *Ca*. B. caroliniensis, absent in Phase I, was also detected through FISH analysis in Phase II (Fig 4). Batch activity tests conducted during Phase II also showed a decline in the nitrite-dependent ammonium removal rate from 1352 mg N g MLVSS^−1^ day^−1^ prior to nitrite depletion (Phase I) to 681 mg N g MLVSS^−1^ day^−1^ after nitrite depletion (Phase II). Nevertheless, the overall activity in R2 was still higher than that in R1 even with a specific nitrite-dependent ammonium removal rate of 575 mg N g MLVSS^−1^ day^−1^ (Fig 5). Under normal operation (Phase I), the NO-dependent ammonium oxidation in the absence of nitrite in both the reactors was insignificant, further supporting the hypothesis that nitrite rather than NO is the preferred electron acceptor and ammonium removal cannot be achieved by *Ca*. B. sinica via direct coupling to NO reduction. In contrast, the ammonium oxidation rates with NO in the absence of nitrite increased more than five times in R2 at 440 mg N g MLVSS^−1^ day^−1^ after nitrite depletion in Phase II compared to 33 and 80 mg N g MLVSS^−1^ day^−1^ in R1 and R2, respectively in Phase I under normal operation (Fig 5). This suggests the selection of AnAOB species capable of utilizing externally supplied NO to oxidize ammonium.

**Figure 5.**
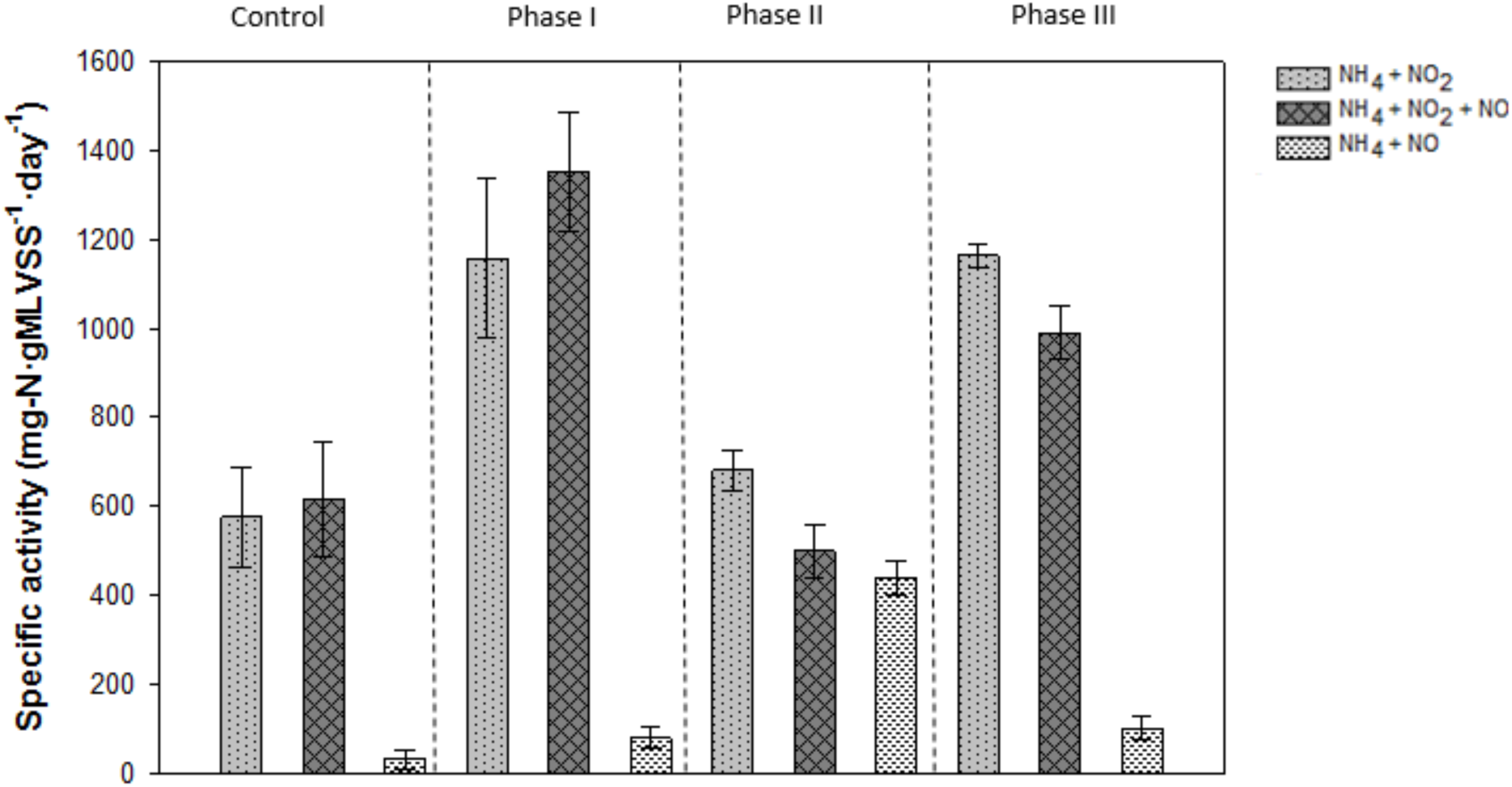
Batch activity experiments with (i) NH_4_ + NO_2_, (ii) NH_4_ + NO_2_ + NO, (iii) NH_4_ + NO in control reactor R1 and experimental reactor R2 a) before nitrite limitation (phase I), b) under nitrite limitation (phase II) and c) after nitrite repletion (phase III)

At the start of experimental Phase III, *Ca*. B. caroliniensis was found to be in higher relative abundance than *Ca*. B. sinica in biofilms forming on the wall of the reactor (Fig 4). *Ca*. Jettenia also showed a recovery, albeit at low abundance (observed in wall samples) in the absence of nitrite (Fig 4). While it cannot be confirmed whether this was a consequence of nitrite depletion in Phase II, the relative abundances of *Ca*. B. caroliniensis and *Ca*. Jettenia in the wall biomass were higher than under normal operation (Phase I in Fig 6). A clear reversal from *Ca*. B. caroliniensis to *Ca*. B. sinica was observed once nitrite was reintroduced between 650 and 680 days (Fig 4). Similar to that detected in Phase II, the increase in relative abundance of *Ca*. B. sinica coincided with the recovery of nitrite-dependent ammonium oxidizing activity of 1163 mg N g MLVSS^−1^ day^−1^ which is comparable to that in Phase I (1352 mg N g MLVSS^−1^ day^−1^) (Fig 4). In addition, the NO-dependent ammonium oxidizing activity also decreased from 440 (Phase II) to 102 (Phase III) mg N g MLVSS^−1^ day^−1^ (Fig 5), further suggesting that increased NO consumption is likely linked to the emergence of *Ca*. B. caroliniensis or *Ca*. Jettenia or both. An extended period under nitrite limitation could have further enhanced the recovery of *Ca*. Jettenia and *Ca*. B. caroliniensis to outcompete *Ca*. B. sinica. Nevertheless, a clear link between nitrite and NO, species selection and their preferred growth mode was demonstrated in this part of the study.

**Figure 6.**
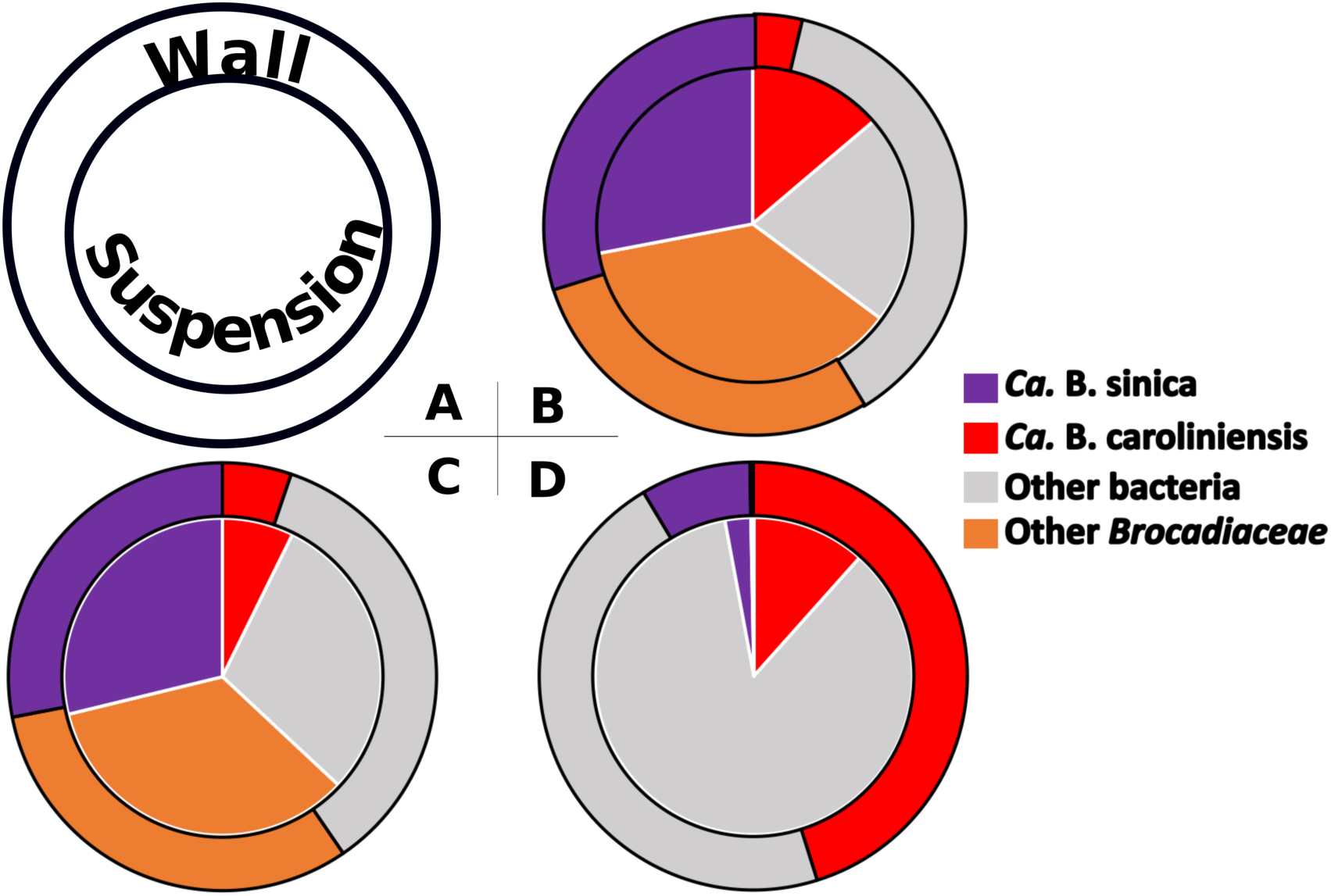
Distribution of dominant OTUs affiliated to *Ca*. B. sinica and *Ca*. B. caroliniensis, along with other AnAOB annotated to *Brocadiaceae* and non-AnAOB on the wall (outer layer) and in suspension (inner core) of (A) <u>R1</u> at day 289 (B), <u>R2</u> at day 265 (C) and <u>R3</u> at day 266 (D). The detailed microbial community composition can be found in Figure S3, Supporting Information.

### Prevailing AnAOB taxa exhibit different biomass morphologies

The predominant AnAOB also displayed a distinct preference for attached growth under the various enrichment conditions. When stable enrichment was achieved, AnAOB biomass was present mainly as suspended granules in R1 and R2 while biofilms attached to the reactor surface were predominantly observed in R3 (Fig S2). When biofilm was transferred from the wall of the reactor into suspension (as indicated by dotted line in Fig 1A – C), a greater increase in the MLVSS of R3 (around 1.5 g L^−1^ at day 230) was observed compared to the other two reactors (less than 0.3 g L^−1^ at days 160, 209, and 258 in R1 and 153, 216, and 253 in R2), suggesting more attached biofilm growth in the primary effluent-fed reactor (R3, Fig 1C). In addition, when nitrite was reintroduced into R2 during experimental Phase III, *Ca*. B. sinica showed a downward trend, more evident in the biofilm samples collected from the walls of the reactor (Fig 4) than in suspension (***p***=0.038). The difference in relative abundance between wall and suspension samples indicates a preference of *Ca*. B. caroliniensis for attached growth. There was no significant difference between AnAOB populations of biomass samples collected from the wall and suspension in R1 and R2, with *Ca*. B. sinica as the dominant AnAOB. However, the relative abundance of predominant *Ca*. B. caroliniensis was four times higher in the biomass collected from the wall than in suspension for R3 (Fig 6), further indicating that this species has a tendency towards attached growth (Fig 6). While granules were found to form in R1 and R2 (Fig 3A and B), with an average particle size of 1.52 and 1527.9 ± 0.078 µm (Table S2) respectively, the morphology of aggregates in R3 was more floc-like (Fig 3C), with a particle size of 310.4 ± 0.2 µm (Table S2). Therefore, it appears that the *Ca*. B. caroliniensis enriched under domestic wastewater conditions containing organic carbon, cannot support granule maturation and instead forms biofilms on walls, in contrast to *Ca*. B. sinica which was mainly detected in the granular biomass.

### Distinct heterotrophic communities coevolved with specific AnAOB taxa

The various enrichment strategies also led to discrete non-AnAOB communities in all three reactors, with higher abundances of non-AnAOB OTUs in R3 compared to R1 and R2 (Figs S1G, H & I). Supplementation of carbon source increased richness (Table S1) in microbial community in R3 (Chao1: 614±42), but selectively enriched for non-AnAOB community (Simpson 1-D: 0.04 ± 0.01) rather than AnAOB (Simpson 1-D: 0.75±0.17). Lower richness was observed when nitrite and ammonium were used as main substrates in R1 and R2 (Chao1: 233 ± 31 and 249 ± 32 respectively). Supplementation of NO led to slightly higher richness but lower evenness in R2 than R1. *Ca*. B. sinica correlated positively with OTUs affiliated to unclassified *Fimbriimonadia* (phylum Armatimonadetes) and unclassified *Anaerolineaceae* (phylum Chloroflexi) in R1 (Spearman’s rho >0.8, ***p***<0.001). While the relative abundance of non-AnAOB was reduced in the presence of NO, the same correlation was nonetheless observed in R2. However, in R3, the non-AnAOB community was supplanted by OTUs affiliated to unclassified *Comamonadaceae* (phylum Proteobacteria) and unclassified Bacteroidetes and a negative correlation was observed between unclassified Bacteroidetes and *Ca*. B. caroliniensis (Spearman’s rho: −0.7, ***p***<0.001). Notably, taxa affiliated to *Comamonadaceae* family were nearly absent in R1 and R2, while they remained the dominant heterotrophic community in biomass both attached to the reactor wall and in suspension in R3, perhaps suggesting that these taxa have metabolic interactions with *Ca*. B. caroliniensis and play a role in biofilm formation (Fig S3).

## DISCUSSION

### N load and organic carbon availability drives AnAOB species selection

Describing factors that drive AnAOB niche differentiation enables populations with desired physiological and growth properties to enhance process control and operational stability to be selected. AnAOB OTUs associated with *Ca*. Brocadia, *Ca*. Kuenenia, *Ca*. Anammoxoglobus, and Ca. Jettenia OTU have all been detected in waste water treatment systems **(3)**. In this study, we systematically tested a range of conditions in order to increase the diversity of AnAOB populations selected from activated sludge, which is commonly used to inoculate both full-scale and lab-scale reactors. This involved providing different metabolites (i.e. ammonium, nitrite, organic carbon and NO) at varying concentrations and loads relevant to mainstream and sidestream PN/A systems. We have demonstrated the reproducible enrichment of two key AnAOB species, *Ca*. B. caroliniensis and *Ca*. B. sinica, from a single seed activated sludge from the tropics and described their preferred ecological niches. *Ca*. B. caroliniensis predominated in all reactors at N loading <750 mg N L^−1^ day^−1^; an increase in N load resulted in a succession towards *Ca*. B. sinica. *Ca*. B. caroliniensis was also the most competitive species in the presence of organic carbon as shown by their prevalence in R3 under conditions of low N loading.

It is possible that *Ca*. B. caroliniensis and *Ca*. B. sinica developed their own niche due to differences in susceptibility to nitrite inhibition since nitrite concentration was increased to increase the N load. A shift in population from *Ca*. B. caroliniensis to *Ca*. B. sinica was consistently observed with increasing nitrite concentration beyond 340 mg NO_2_-N L^−1^ in the feed (i.e., 170 mg NO_2_-N L^−1^ in the reactor). This is within the range of inhibitory concentrations reported for AnAOB of 40 mg N L^−1^ to 400 mg N L^−1^ **(16-18, 29-31)**. However, this has not been specifically investigated at species level.

An alternative explanation could be that *Ca*. B. caroliniensis and *Ca*. B. sinica have different intrinsic kinetic properties. Using enriched planktonic cells, the nitrite affinity constant and specific growth rate of *Ca*. B. sinica were determined to be 0.47 mg N L^−1^ and 0.33 day^−1^ (corresponding doubling time of 2.1 days) **(28, 32)**, which is the highest maximum growth rate ever reported for AnAOB. This indicates that *Ca*. B. sinica are r-strategists and would grow at high N loading rates **(32)** as observed in our study. However, a similar estimation for *Ca*. B. caroliniensis is missing. Metagenomic analysis revealed that *Ca*. B. caroliniensis have multiple copies of nitrite/formate transporters (*focA*) that provide a competitive advantage at low nitrite concentrations due to a low intrinsic nitrite affinity constant **(14)**. This could potentially help them scavenge nitrite from heterotrophic denitrifiers in the presence of organic carbon.

Despite the wide range of environmental conditions applied across the three enrichment reactors, *Ca*. Brocadia remained the most dominant phylotype throughout the enrichment process, while *Ca*. Kuenenia and *Ca*. Anammoxoglobus commonly found in engineered systems were not detected. It is possible that *Ca*. Kuenenia and *Ca*. Anammoxoglobus were not present in the inoculum. Therefore, it is possible that different species may emerge with the conditions tested in this study depending on the AnAOB diversity in the inoculum **(33)**.

### NO can select for specific AnAOB taxa

NO was provided in R2 to exert oxidative stress and also to potentially select for NO-utilizing AnAOB **(34)**. While the presence of NO did not appear to suppress the growth of AnAOB, the presence of NO provided a competitive advantage to *Ca*. Jettenia in R2 to a maximum relative abundance of 23% along with *Ca*. B. caroliniensis at low N loading (Fig 1). Little is known about the ecological and metabolic drivers of the niche of *Ca*. Jettenia, probably because they are generally less abundant than other genera of AnAOB **(3)**. Low nitrite concentrations were shown to encourage the proliferation of *Ca*. Jettenia over *Ca*. B. sinica **(4)**, consistent with this study. However, *Ca*. Jettenia was much less abundant than *Ca*. B. caroliniensis at low nitrite loading rates. This suggests their growth was limited by the low NO availability compared to nitrite. In addition, *Ca*. Jettenia might also prefer a much lower N load as shown by the decrease in relative abundance at higher N loading rate compared to *Ca*. B. caroliniensis. Nevertheless, we show that two phylogenetically distant AnAOB species, *Ca*. B. caroliniensis and *Ca*. Jettenia, can coexist in the same system.

The coevolution of *Ca*. B. caroliniensis and *Ca*. Jettenia might also suggest that *Ca*. B. caroliniensis could utilize NO. A similar nitrite reduction pathway in *Ca*. B. caroliniensis and *Ca*. Jettenia was suggested after a *nirK* homologue was detected from the metagenomic analysis of *Ca*. B. caroliniensis. **(14, 35)**. The significant increase in NO-dependent ammonium oxidation following nitrite limitation was concomitant with the recovery of *Ca*. B. caroliniensis and *Ca*. Jettenia populations. It is conceivable that *Ca*. B. caroliniensis utilize a *nirK* homologue or a novel nitrite reductase employing the conventional NO-dependent pathway for hydrazine production or possess the ability to switch to a NO-dependent pathway in the absence of nitrite. An alternate pathway for NO production through oxidation of hydroxylamine by a hydroxylamine oxidoreductase (*hao*)-like protein was indeed detected by Park et al. **(14)** and proposed that this alternate pathway could potentially be activated under nitrite limitation. NO was also shown to oxidize ammonium to dinitrogen gas under nitrite limitation in *Ca*. B. fulgida **(21)** and canonical *nirS* was absent in its genome **(36)**, suggesting the involvement of a yet-to-be identified nitrite reductase in *Ca*. Brocadia species for reduction of nitrite.

In contrast, when nitrite was present at significantly higher concentrations than NO, NO-dependent ammonium oxidation was negligible, further corroborating the assertion that nitrite rather than NO was the preferred electron acceptor and that ammonium removal cannot be achieved by *Ca*. B. sinica via direct coupling to NO reduction (Fig 5). While *Ca*. B. sinica was not completely inhibited in the absence of nitrite, its relative abundance decreased under nitrite limitation as discussed earlier. This supports the study by Oshiki et al. **(37)** demonstrating that *Ca*. B. sinica does not utilize NO and ammonium for hydrazine synthesis, but instead uses hydroxylamine and ammonium.

### The predominant AnAOB species determine biomass retention strategy

High retention of AnAOB in reactors is a crucial factor for optimal operation due to the slow growth rate of these bacteria. This can be achieved through biofilm attachment to carriers, formation of granular biomass aggregate and other separation techniques such as membrane filtration to prevent washout of AnAOB. The choice of biomass retention in Anammox systems may be guided by the growth mode of prevailing AnAOB species and coexisting microbial community under specific operational conditions. In this study, we demonstrated that the AnAOB community and their aggregation states might be distinct under mainstream and sidestream conditions. *Ca*. B. caroliniensis, likely persisting under mainstream conditions, exhibited a preference for attached biofilm growth. High diversity and abundance of heterotrophic species in R3 was also observed in the presence of complex organic carbon. Particularly, *Comamonadaceae* remained one of the most abundant heterotrophs in the system both in suspension and in the biofilm. *Comamonadaceae* bacterium are commonly found in biofilm forming communities **(38, 39)** suggesting their potential role in assisting biofilm formation. Attached growth of *Ca*. B. caroliniensis was also observed in a full-scale process treating anaerobic digester liquid supplied with glycerol as the external carbon source **(14)**. In this instance, the use of carriers can be used to provide a large surface area to achieve high biomass retention **(40)**. Carriers supporting attached biofilm growth can be applied in various configurations, for instance, in rotating biological contactors **(41)**, moving bed biofilm reactors **(42, 43)** and sequencing batch biofilm reactors **(44)**. However, when AnAOB form granules in a *Ca*. B. sinica dominated system (as observed in R1 and R2 operated under side stream conditions), physical separation of dense Anammox granules can be achieved by using a hydrocyclone to separate slow growing Anammox bacteria from incoming solids as used in DEMON SBR systems **(45)** or granular systems with lamella separators for granule retention **(46)** or integrated fixed film activated sludge (IFAS) configurations with a settler **(47)**. Despite the absence of externally supplied organic carbon, heterotrophs belonging to class *Fimbriimonadia* (phylum Armatimonadetes) and family *Anaerolineaceae* (phylum Chloroflexi) proliferated in R1 and R2 to a relative abundance of 10-15% in. Gao et al. **(48)** suggested an important role of *Anaerolineaceae* as cores or carriers for granule formation in Anammox sludge and their increase in abundance over time would suggest that they may have supported granulation in both R1 and R2. However, the role of heterotrophic bacteria and their interaction with AnAOB cannot be fully uncovered in this study and will require further investigation.

## CONCLUSIONS

- Multiple AnAOB species can be enriched from activated sludge. *Ca*. B. caroliniensis dominated at low N loads, both in the presence and absence of organic carbon and under nitrite limitation, suggesting that it has a high metabolic versatility, and is therefore more competitive and persistent in mainstream systems. Their tendency to form attached biofilms suggests that carrier systems would be a more suitable strategy for biomass retention.
- *Ca*. B. caroliniensis is replaced by *Ca*. B. sinica that form granules at higher N load, which would dominate in side stream systems devoid of organic carbon.
- Although *Ca*. Jettenia could grow in the presence of externally supplied NO, their disappearance in environments rich in nitrite suggests that they are not competitive in wastewater treatment systems.
- Collectively, this study provides insight into understanding the relationship of species selection, growth morphology and process conditions in mainstream and sidestream applications and can have important implications in process design, control and management of the Anammox process at the species level in full scale waste water treatment systems.

## MATERIALS AND METHODS

### Laboratory-scale sequencing batch reactor set up and operation for AnAOB enrichment

Three sequencing batch reactors (SBRs), each with a working volume of 4 L, were seeded with activated sludge from a full-scale water reclamation plant (WRP) in Singapore. R1 and R2 were fed with synthetic medium devoid of organic carbon while R3 received primary effluent collected from the WRP once a week comprising complex organic carbon source. Synthetic medium was prepared as (g L^−1^): KHCO_3_ 1.25, KH_2_PO_4_ 0.025, CaCl_2_·6H_2_O 0.3, MgSO_4_·7H_2_O 0.2 and FeSO_4_·7H_2_O 0.025 with gradual increase in ammonium (from 30 to 280 mg N L^−1^) and nitrite concentration (39 to 350 mg N L^−1^) and 1.25 ml L^−1^ of trace mineral solution as described by van de Graaf et al. (1996). Argon/CO_2_ (95/5%) was sparged continuously at 25 mL min^−1^ throughout the anoxic phase to prevent ingress of oxygen in R1 whereas R2 was sparged with both Argon/CO_2_ and NO with a combined sparging rate of 25 mL min^−1^ to a final NO gas phase concentration of 400 ppmv to impose oxidative stress. The selected concentration of 400 ppmv was lower than the previously reported tolerance threshold of 600 ppmv for AnAOB **(22)**. R1 and R2 were operated in cycles of 12 h, each cycle comprised of 5 min of feeding, 108 min of anoxic cycle, 67 min of settling and decanting. Initial ammonium and nitrite concentrations were maintained at 20 mg N L^−1^ within the reactor for the first seven weeks resulting in a hydraulic retention time (HRT) of 24 h (two litres of synthetic waste water were fed into the reactor each cycle). Upon achievement of 100 % ammonium removal, NH_4_^+^ and NO_2_^−^concentrations in the feed were then increased in steps of 20 mg N L^−1^. Influent NO_2_^−^: NH_4_^+^ was maintained at a molar ratio of 1.3 close to theoretical stoichiometry **(49)**. HRT was gradually decreased from 24 h to 16 h for R1 and from 24 h to 12 h for R2, in accordance with the N removal capacity. Thus, R2 was operated at a higher N loading rate compared to R1 due to shorter cycle time concomitant with higher N removal rates in R2. The pH was not controlled in either R1 or R2 and varied between 7.2 and 7.8.

R3 was operated in cycles of 8 to 12 h, each cycle comprised of 2 h feeding, 5 to 9 h anoxic phase (depending on the cycle length applied), and 1 h of settling and decanting. Before settling, the reactor was sparged with Argon/CO_2_ for 5 min to strip out the nitrogen gas produced during the anoxic phase to improve sludge settle-ability. In each feeding period, 2 L of primary effluent supplemented with nitrite was added, resulting in a HRT of 16 to 24 h. Nitrite was adjusted according to the ammonium concentration at a molar ratio of 2:1 and stored in a chiller at 4 °C to minimize degradation. The nutrient composition of primary effluent was measured after addition of nitrite with average values shown in Table S1. Slow feeding of 2 h was applied to minimise oxygen introduction and temperature shock from primary effluent stored in the chiller. The pH of the reactor was not controlled and varied between 7.6 and 8.5 due to denitrification activity. For enrichment purpose, the SRT was not controlled in all three reactors whereby sludge loss only occurred through sampling for nutrient and solids analyses.

A heating jacket was connected to maintain the SBR at 35 ± 0.05°C for R1 & R2 and 33 ± 1°C for R3. Dissolved oxygen (DO) concentration and pH were continuously monitored using Mettler Toledo InPro6050 DO sensor and Mettler Toledo-InPro 3250i pH sensor, respectively. Samples were collected periodically at the end of the cycle and filtered immediately with 0.2 µm filters for nutrient analyses. Mixed liquor samples were collected in the middle of the anoxic phase for DNA extraction. To determine the microbial composition of the biofilm on the reactor surface, biomass samples from the wall was collected from three random locations after draining the reactor at day 289 for R1, day 265 for R2 and 266 for R3. Both collected suspended and biofilm samples were snap-frozen in liquid nitrogen and stored at −80 °C until extraction. Biofilm on the surface of the reactor was periodically cleaned to determine total mixed liquor suspended solids (MLSS) and the volatile fraction (MLVSS) and the proportion of biomass that was in suspension versus attached growth. Suspended biomass samples were also collected for particle size analysis, and light microscopy imaging after stable enrichment was attained.

### NO depletion and repletion experiment

To further validate the effect of NO as a selection pressure for AnAOB species selection, R2 was subjected to gradual nitrite depletion and repletion while maintaining availability of NO as an electron acceptor across three experimental phases after stable operation was attained: **Phase I -**normal operation prior to nitrite depletion, ammonium and nitrite concentration in the feed were 280 and 350 mgN L^−1^, respectively with continuous supply of NO at 400 ppmv in the gas phase (before day 563); **Phase II**-nitrite limited operation (day 564-640) whereby nitrite was reduced stepwise from 50 to 0 mg NO_2_-N L^−1^ while ammonium and NO were maintained at 50 mg NH_4_-N L^−1^ and 400 ppmv, respectively; **Phase III** (day 640-687) nitrite was gradually reintroduced from 0 to 70 mg NO_2_-N L^−1^ with the aforementioned ammonium and NO concentrations in Phase II. Biomass samples were collected twice a week from the mixed liquor throughout the experiment however wall biomass samples were only collected from Phase III due to the limited amount of wall biomass. At each experimental phase, batch activity tests were conducted in triplicate with 80 mg NH_4_^+^-N L^−1^, 100 mg NO_2_^−^-N L^−1^ and/or 400 ppmv NO in gas phase under the following conditions with (i) ammonium and nitrite only in the absence of NO, (ii) ammonium, nitrite and NO, and (iii) presence of ammonium and NO only in the absence of nitrite. The Anammox activity of R1 under normal operation with ammonium and nitrite supplied as substrate served as the control. In all batch activity tests, mixed liquor samples were collected every 30 min and filtered through 0.22 µm Milipore filters for nutrient analysis.

### Chemical analysis

All samples collected for nutrient analysis were measured for ammonium, nitrite and nitrate. Ammonium was measured using Hach® kits, nitrate and nitrite were analyzed using ion chromatography (Prominence, Shimadzu). MLSS and MLVSS were analyzed according to the standard methods (Eaton et al. 2005). NO was measured in the gas phase using an online chemiluminescence analyzer (Model: 42i, Thermoscientific). Particle size analysis was carried out using laser diffraction particle size analyzer (Model: SALD-MS30, Shimadzu).

Suspended biomass was collected from each reactor and subject to size analysis. 1 mL biomass was dispersed on the surface of petri dish, and images were taken by AxioObserver Z1 inverted epifluorescent microscope (Leica, Germany) with brick/seal function. Images were then analyzed with image J **(50)** Analyze particles function.

### Microbial community profiling

Genomic DNA was extracted from biomass samples using FastDNA^™^ SPIN Kit for Soil (MP Biomedicals, USA) with optimization according to Albertsen et al. (2015). Pair-end 16S rRNA amplicon sequencing was conducted by DNAsense (http://dnasense.com/) at the Aalborg University (Denmark) with primer set 515F (5’-GTGCCAGCMGCCGCGGTAA-3’) and 806R (5’-GGACTACHVGGGTWTCTAAT-3’) (Caporaso et al. 2011) by Illumina Miseq platform as described in Law et al. (2016). Detailed data analysis can be found in supporting information (SI).

### Clone library construction and phylogenetic tree analysis

To further confirm the species level taxonomy identification, four 16S rRNA gene clone libraries were constructed from samples collected on days 74, 280 from R1 and 85, 270 from R2. Sequences obtained from clone library and 16S rRNA amplicon sequencing were used to generate phylogenetic tree by ARB. Methods for clone library and phylogenetic tree construction are described in detail in SI.

### Fluorescent *in situ* hybridization

Suspended biomass samples were collected and fixed with 4% PFA overnight. After washing with 1 × PBS (130 mM sodium chloride, 10 mM sodium phosphate buffer, pH 7.2), the biomass samples were stored with 1:1 100% ethanol:1x PBS at −20°C. FISH was performed on crushed biomass samples according to the method described by Daims et al. **(51)** with probes listed in Table 1. The slides were viewed using a Zeiss LSM 780 inverted confocal microscope (Carl Zeiss, Jena, Germany).

**Table 1.**
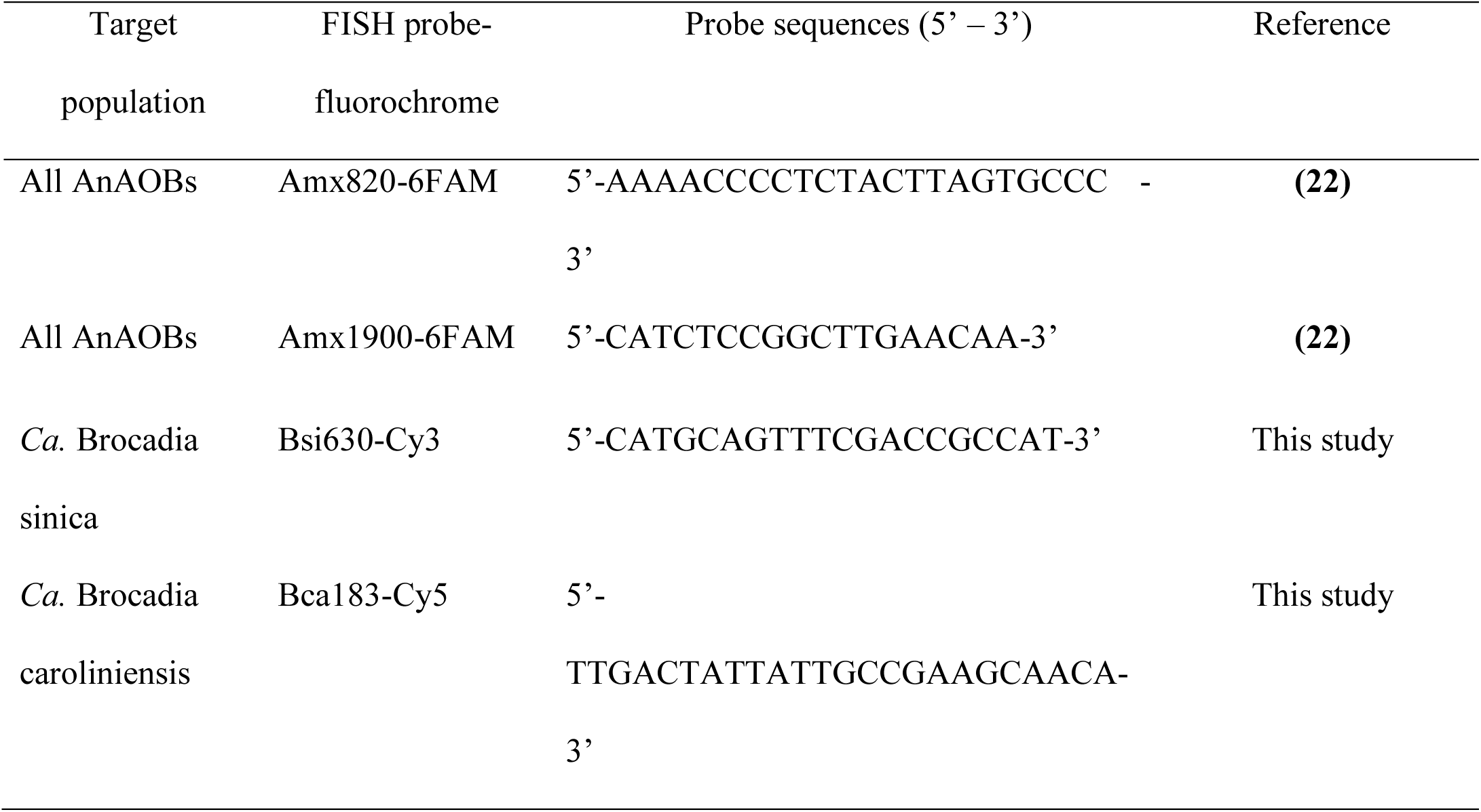
List of rRNA-targeted oligonucleotide probes used for FISH analysis.

## Data availability

All raw16S rRNA amplicon sequences used in this manuscript are available in NCBI under Bioproject PRJNA604076.

## ACKNOWLEDGEMENTS

This research was supported by the Singapore National Research Foundation and Ministry of Education under the Research Centre of Excellence Programme, and by a program grant from the National Research Foundation (NRF), project number 1301-IRIS-59. We would like to acknowledge the assistance of Mr. Larry Liew and staff from the Public Utility Board (PUB) of Singapore for the weekly collection of primary effluent and Mr. Eganathan Kaliyamoorthy for his help in conducting solids analysis.

